# TCF3::HLF Orchestrates an Enhancer-Promoter Network with Activation of MEF2C to Promote Immature HSC gene Expression in Leukemia

**DOI:** 10.1101/2024.11.26.625006

**Authors:** Valdemar Priebe, Aneta Drakul, Bartimée Galvan, Júlia Aguadé-Gorgorió, Hanna K. A. Mikkola, Beat Bornhauser, Raffaella Santoro, Jean-Pierre Bourquin

**Affiliations:** Division of Oncology and Children’s Research Centre, University Children’s Hospital Zurich, Zurich, Switzerland; Department of Molecular Mechanism of Disease, University of Zurich, Zurich, Switzerland; Department of Molecular, Cell and Developmental Biology, University of California Los Angeles, Los Angeles, CA, USA; Eli and Edythe Broad Center for Regenerative Medicine and Stem Cell Research, University of California Los Angeles, Los Angeles, CA, USA; Jonsson Comprehensive Cancer Center, University of California Los Angeles, Los Angeles, CA, USA; Molecular Biology Institute, University of California Los Angeles, Los Angeles, CA, USA

## Abstract

Oncogenic fusion transcription factors (TFs) frequently drive hematopoietic malignancies by altering gene expression in key developmental programs. TCF3::HLF is a fusion TF that characterizes a rare, treatment-resistant subtype of B-cell acute lymphoblastic leukemia (t(17;19) TCF3::HLF-positive B-ALL). Despite its clinical significance, the mechanisms by which TCF3::HLF induces leukemia are unclear. We used HiChIP mapping and genetic interference to analyze TCF3::HLF at the 3D-genome level, revealing enhancer-promoter interactions that control gene activation or repression. Notably, TCF3::HLF directly regulates *MEF2C* expression through its enhancer, as interference disrupted *MEF2C* transcription and inhibited leukemia propagation. This disruption also diminished embryonal hematopoietic stem cell (HSC) gene signatures and restored mature HSC and B-lymphoid markers. These findings highlight *MEF2C* as a critical component of the transcriptional network reprogrammed by TCF3::HLF. Our study provides insight into how TCF3::HLF rewires the 3D genome to drive leukemia and serves as a resource for further exploration of the TCF3::HLF regulome.

**Teaser:** Unraveling the 3D genomic interactions mediated by TCF3::HLF fusion protein in t(17;19) positive acute lymphoblastic leukemia.

## Introduction

Studies of the three dimensional (3D) genome have provided relevant insights into gene regulation that cannot be described in direct linear fashion, such as enhancer-promoter (E-P) contacts (*1–3*). Increasing evidence shows that enhancer malfunction is a key process that drives the aberrant regulation of oncogenes in cancer. Differential genome organization between healthy and malignant cells has been observed and can amongst other be attributed to chromosomal translocations leading to both alterations of genome structure or hijacking of enhancers by oncogenic fusion proteins (*4*, *5*).

Chromosomal translocations resulting in oncogenic fusion of transcription factors (TFs) involving hematopoietic master regulators are frequent genetic abnormalities in acute leukemias (*6–8*). These chimeric TFs may take over the functions of master regulators that typically control gene expression programs defining cellular identity by hijacking enhancer elements in cooperation with other TFs (*8–10*). These oncogenic TFs disrupt fundamental cellular processes, including self-renewal, differentiation, and proliferation, thereby imparting leukemogenic potential to hematopoietic stem and progenitor cells (HSPCs) (*11*).

The chromosomal translocation t(17;19)(q22;p13) generates the fusion protein TCF3::HLF that defines a rare subtype of B-cell acute lymphoblastic leukemia (t(17;19) TCF3::HLF positive ALL) (*12–15*). This chimera consists of the transactivation domains of TCF3, a TF that drives lymphoid development, fused to the DNA-binding and dimerization domains of the leukemia-associated TF hepatic leukemia factor (HLF), which is a regulator of multipotent hematopoietic progenitors and is rapidly downregulated upon differentiation (*14*, *16* – *19*). Accordingly, the resulting chimeric TCF3::HLF TF rewires the transcriptional programs towards a stem-like/myeloid expression lineage profile and promotes leukemogenesis in cells of committed lymphoid origin (*20*, *21*). t(17;19) positive ALL has less than 1% prevalence in B-ALL pediatric cases. Although inclusion of the BCL2 inhibitor venetoclax and of CD19 directed immunotherapy resulted in sustained remissions (*21*, *22*), TCF3::HLF positive ALL still remains one of the most resistant ALL subtypes.

Previous chromatin immunoprecipitation (ChIPseq) analyses of TCF3::HLF in t(17;19) positive ALL cells showed preferential association of TCF3::HLF with enhancer sequences (*20*). However, knowledge about the gene regulatory circuitry mediated by TCF3::HLF remains limited given the difficulty to predict distant interactions between regulatory elements in the genome. To capture the relevant clusters of direct TCF3::HLF target genes, we performed TCF3::HLF-HiChIP to identify the 3D-architecture of TCF3::HLF bound E-P interactions that is underlying the oncogenic activity in t(17;19) positive ALL. By integrating TCF3::HLF spatial genomic information with chromatin and transcriptomic data, we identified networks of activating and repressing E-P interactions that are directly driven by TCF3::HLF. This approach identified previously undetected TCF3::HLF target genes that are specifically upregulated in t(17;19) positive ALL relative to other ALL types and implicated in cell-cell interaction or neuronal development. The most prominent functional TCF3::HLF-bound E-P interaction was with the myocyte enhancer factor 2C (*MEF2C*). We demonstrated that *MEF2C* depends on the interaction of TCF3::HLF with the enhancer and showed that it controls significant portions of the TCF3::HLF transcriptional program, mirroring fetal hematopoietic stem cell features with functional essentiality for the leukemia. These findings represent a paradigm for how oncogenic fusion TFs can rewire the 3D-genome to drive malignancy.

## Results

### A map of TCF3::HLF 3D-genomic interactions in t(17;19) positive ALL cells

To map the 3D-genomic interactions associated with the fusion protein TCF3::HLF in t(17;19) ALL cells, we performed HiChIP, a method that combines chromatin immunoprecipitation (ChIP) and chromosome conformation capture (HiC) to directly capture long-range DNA interactions associated with a protein of interest (*23*). We generated TCF3::HLF-HiChIP maps from two independent replicates using the TCF3::HLF positive cell line HAL-01 (*12*) whose TCF3 wild-type allele was knocked out (*20*), thereby enabling selective immunoprecipitation of the fusion protein when using antibodies against TCF3 (**Fig. 1A**).

**Fig. 1.**
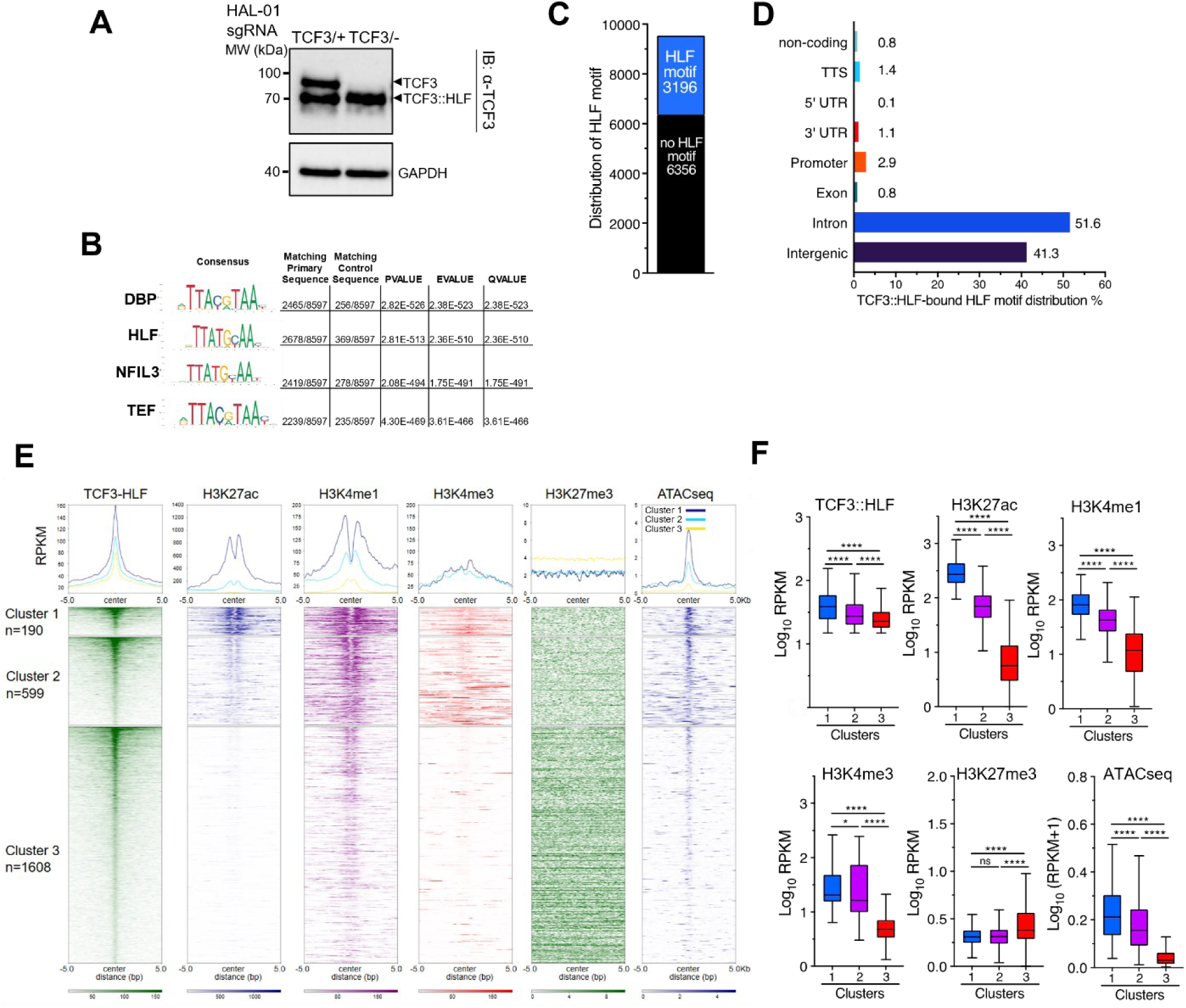
TCF3::HLF 3D-genomic interactions in t(17;19) positive ALL cells. (**A**) Anti-TCF3 immunoblot of HAL-01 cells expressing TCF3::HLF. Endogenous TCF3 was knocked out by CRISPR editing using sgRNA. Expression of TCF3 and TCF3::HLF are shown, GAPDH serves as loading control. (**B**) Motif enrichment of HiChIP-detected TCF3::HLF binding sites (n=9552). (**C**) Distribution of HiChIP-detected TCF3::HLF binding sites containing HLF motif. (**D**) Genomic annotation of HiChIP-detected TCF3::HLF binding sites containing HLF motif. (**E**) Heatmaps of TCF3::HLF, H3K27ac, H3K4me1, H3K4me3, and H3K27me3 occupancy and chromatin accessibility by ATACseq (*74*) at HLF motif of parental HAL-01 cells. Sequences were clustered in 3 groups based on the histone modification content. (**F**) Boxplots showing the average RPKM read density of regions centered on HLF motifs. Statistical significance (P-values) was calculated with two-tailed T test (*<0.05; ****< 0.0001, ns: not significant).

We identified more than 49’000 significant interaction pairs (FDR <0.01) distributed across the 23 human chromosome pairs (**fig. S1, A and B**). These interaction pairs comprised 9’552 unique sequences that exhibit a significant enrichment in the DNA recognition motif of basic leucine zipper (*bZIP*) TFs, including HLF that is recognized by TCF3::HLF (*P* 2.81×10^-513^) (**Fig. 1, B and C**). Among these sequences, 3’196 (33%) contained an HLF binding motif, underscoring the specificity of TCF3::HLF-HiChIP (**Fig. 1C**). The TCF3::HLF-bound regions with HLF motif were predominantly located at intergenic regions (42%) and introns (52%) but only a small fraction (2.9%) in gene promoter regions (**Fig. 1D**), a result consistent with previous TCF3::HLF ChIPseq data (*20*). Next, we analyzed the chromatin state of these regions with HLF motif by assessing the occupancy of active (H3K27ac, H3K4me3, and H3K4me1) and repressive (H3K27me3) histone modifications, and chromatin accessibility (ATACseq) at sequences containing the HLF binding motif (**Fig. 1E**). We first selected the top 75% TCF3::HLF enriched regions at the HLF motif. As expected, we observed high signal for active histone modifications and chromatin accessibility at TCF3::HLF-bound sites whereas there was no evident enrichment for H3K27me3, indicating that these sequences display active chromatin states. The presence of sharp peak signals for H3K27ac and H3K4me1 relative to H3K4me3 at HLF motif recalls an enhancer chromatin signature. Hierarchical clustering defined three groups according to the levels of the analyzed histone modifications (**Fig. 1F**). Cluster 1 displayed higher enrichment for H3K27ac, H3K4me1, and chromatin accessibility than clusters 2 and 3. Cluster 2 was enriched in H3K4me1 while showing lower H3K27ac content relative to cluster 1, a chromatin signature characterizing primed or poised enhancers (*24*). Finally, cluster 3 was mostly depleted of active histone modifications and contained higher H3K27me3 levels, indicating that these sequences are in a repressive state.

To identify genes directly regulated by TCF3::HLF, we focused on interaction pairs between inter -or intragenic regions containing the HLF motif (I_HLF_) and promoters (P) (*n*=1409) (**Fig. 2A**). Using Rank Ordering of Super-Enhancers (ROSE) (*25*, *26*) and the Fantom5 database (*27*, *28*), we found that a large majority of these I_HLF_-P interactions (72%) contained a signature of enhancer, a result consistent with the analysis of the histone modifications (**Fig. 2B**). By intersecting previously published RNAseq of the parental leukemia cell line HAL-01 upon *TCF3::HLF*-KO (*20*), we found 1057 (75%) I_HLF_-P pairs corresponding to the promoters of genes significantly upregulated (512, adj.p<0.05) or downregulated (545, adj.p<0.05) upon TCF3::HLF loss (from now on named E-P genes, **Fig. 2, C and D, fig. S1B**). We analyzed the distribution of up- and downregulated E-P genes in the three clusters obtained from the analysis of histone modifications at I_HLF_ regions bound by TCF3::HLF (**Fig. 2E**). We found that 67% of E-P genes within cluster 1 were downregulated upon TCF3::HLF loss (**Fig. 2E**), suggesting that TCF3::HLF can positively regulate the expression of these genes through its interaction with active enhancers and 3D-contacts with the corresponding promoters. In contrast, we did not find differences in the distribution of up- and downregulated genes in cluster 2 and 3.

**Fig. 2.**
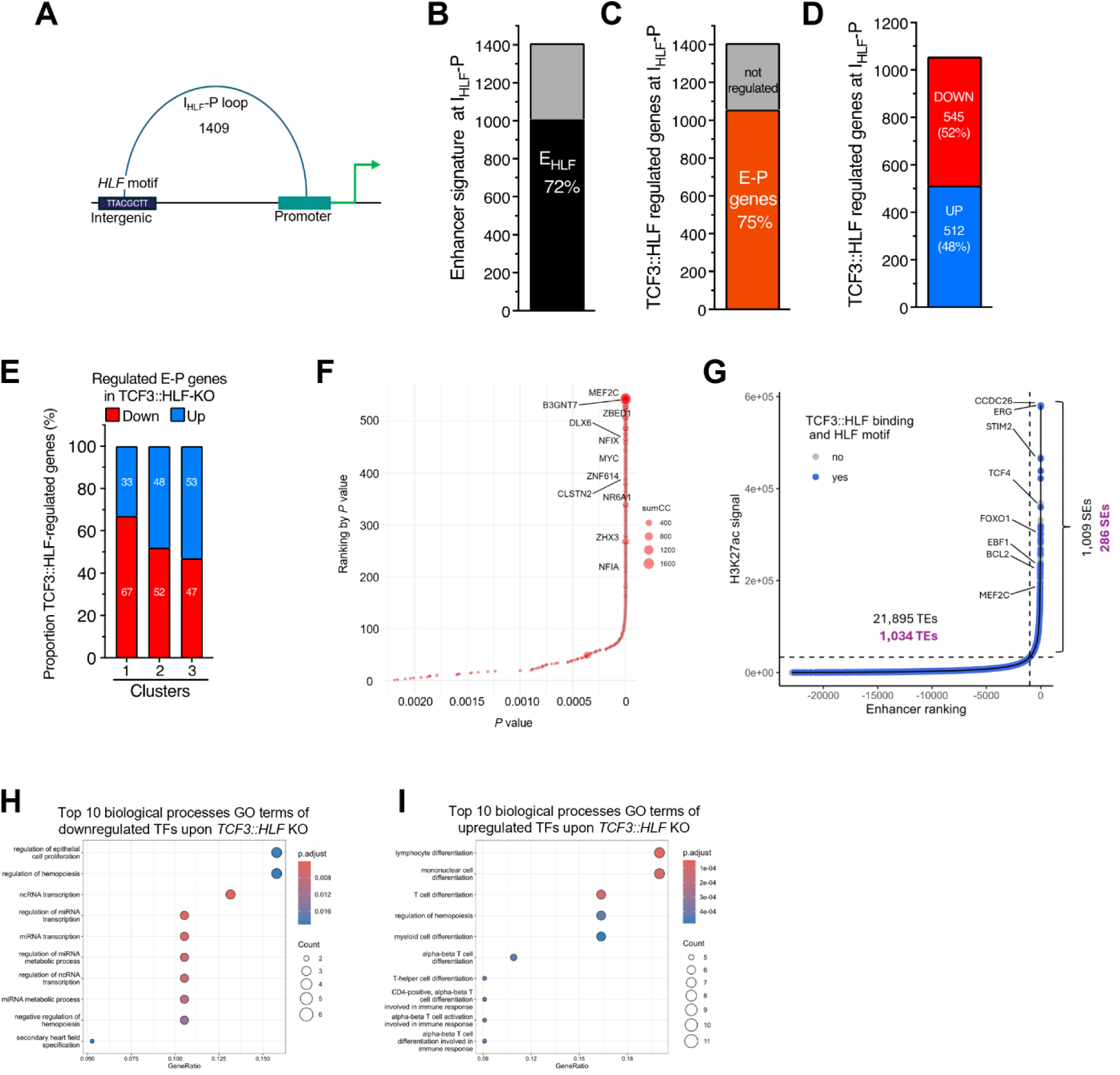
Enhancer-promoter pairs mapped with HiChIP identify TCF3::HLF direct target genes. (**A**) Illustration of interaction pairs between intergenic and promoter regions (I_HLF_-P) that are bound by TCF3::HLF and contain HLF motif. (**B**) Number of I_HLF_-P interaction containing enhancer signature (E_HLF_). (**C**) Number of TCF3::HLF regulated genes at I_HLF_-P pairs (E-P genes). (**D**) Number of significantly upregulated and downregulated E-P genes upon TCF3::HLF-KO. (**E**) Distribution of upregulated and downregulated E-P genes among the three clusters obtained with the histone modification analysis shown in Fig. 1F. (**F**) Plot showing the P value and the ranked order of interaction of I_HLF_-P pairs for the 1057 E-P genes significantly regulated upon TCF3::HLF-KO. The size of the data points indicates the cumulative interaction value (sumCC). (**G**) Plot showing the ranked order of H3K27ac signal using ROSE analysis. Typical enhancers (TE) and super enhancers (SE) are annotated as sites with TCF3::HLF binding (Blue) or not (Gray). Enhancers regulating an E-P gene of interest according to HiChIP are annotated with target gene. (**H, I**) Plots showing gene ontology terms for biological processes of TFs from E-P genes that are downregulated (**H**) and upregulated (**I**) upon TCF3::HLF-KO.

To identify E-P genes with the most significant I_HLF_-P interaction, we first calculated the sum of contact counts (sumCC), a value that represents the cumulative counts of all significant I_HLF_-P interactions and consequentially their contact frequency (**Fig. 2F**). By using this approach, we identified not only the most significant I_HLF_-P interactions but also the ones with highest contact frequency. Among these I_HLF_-P interactions, the most significant was between the promoter of *MEF2C* and the TCF3::HLF-bound intergenic sequence that was recognized as enhancer by the ROSE and FANTOM dataset analysis. MEF2C is a regulator of hematopoietic self-renewal and differentiation, and it has been associated with increased risk of relapse when highly expressed in several subtypes of AML (*29–34*). However, its role in TCF3::HLF positive ALL has not yet been investigated, a task that we set to investigate in this study (see below). Consistent with previous work (*20*), we also detected a *MYC* enhancer-promoter interaction as one of the highest interactions within the E-P genes. Furthermore, among the E-P genes positively regulated by TCF3::HLF, we found *BCL2* (**Fig. 2G**). This result is consistent with recurrent high sensitivity observed with the BCL2-specific BH3-mimetic venetoclax in the clinic, which is now recommended for use in combination chemotherapy in first line for TCF3::HLF positive leukemia (AIOEP-BFM 2017 study, EudraCT 2016-001935-12) (*20*, *21*).

To obtain insight into the possible network of transcriptional regulators that are directly controlled by the TCF3::HLF, we identified 94 TFs among E-P genes that were significantly regulated upon *TCF3::HLF*-KO (39 downregulated and 55 upregulated). Among the 39 TFs that are activated by TCF3::HLF and detected in active clusters 1 and 2 based on chromatin marks, we found MEF2C, MYC, MESP1 and DLX6 genes among others, which regulate proliferation, miRNA metabolism and contextual repression of hematopoiesis (*30*, *33*, *35–39*) (**Fig. 2H**). Among the 55 TF genes that are repressed by the oncogenic fusion, we found HLX, SOX4, FOXP1 and BCL6, which regulate hematopoiesis and myeloid differentiation (*40–45*) (**Fig. 2I**). These results highlight how TCF3::HLF can regulate a network of TFs directly to modulate the cellular state in critical functional hallmarks of cancer such as cell proliferation and self-renewal.

We next explored which of the direct targets that are activated by TCF::HLF would be specifically expressed in TCF3::HLF ALL compared to all other ALL subtypes. We took advantage of a comprehensive dataset containing >10’000 pediatric patients with cancer and long-term survivors (https://www.stjude.cloud) (*46*) to compare the expression of E-P genes among 24 ALL subtypes. We identified 22 E-P genes that are preferentially expressed in TCF3::HLF ALL, providing further candidate biomarkers for this leukemia subtype. Gene ontology analysis for biological processes shows involvement of cell-cell adhesion (**Fig. S2)**. Among the most specifically expressed genes, we detected CLSTN2 and B3GNT7 (**Fig.2F, fig. S1, C and D**). Both genes encode for membrane proteins with possible roles for cell adhesion, invasion, and metastasis, with a role in neuronal development (*47*, *48*). These warrant further functional investigation for possible mechanisms of action and targets for intervention. Taken together, 3D conformational mapping provides new insights into candidate direct targets of TCF3::HLF and the 3D-organization of the TCF3::HLF oncogenic program.

### TCF3::HLF builds a hub of long-range interactions between super enhancer elements and *MYC* promoter to drive MYC expression

The HiChiP analysis of TCF3::HLF occupancy provides new unbiased information on 3D-genome organization within TCF3::HLF-bound regions. We have previously reported that TCF3::HLF associates with the super enhancer 2 (SE2) within the blood enhancer cluster (BENC) to regulate *MYC*, as disruption of the HLF motif in SE2 reduced its interaction with the *MYC* promoter and downregulated *MYC* expression (*20*). HiChIP data not only validated this interaction but also provided additional TCF3::HLF chromatin anchors and binding sites (**Fig. 3A**). First, we observed TCF3::HLF association with *MYC* promoter that was not detectable in previous TCF3::HLF-ChIPseq analyses, supporting a direct role of TCF3::HLF in directly regulating *MYC* expression. Second, we identified additional TCF3::HLF binding sites upstream of and within the BENC locus, thereby constituting a network of interactions involving the *MYC* promoter. One strong TCF3::HLF position was detected upstream and outside of the annotated BENC enhancer sequences (Anchor, **Fig. 3A**). This site appears to act as an anchor to the critical TCF3::HLF binding sites at the BENC and *MYC* promoter. This anchor region is depleted of H3K27ac while enriched in H3K4me1, which is proposed to mark enhancer poised states, and is categorized within cluster 3 in our histone state analysis (*49*, *50*) (**Fig. 1E**, **Fig. 3A**). This anchor site may play an important role in stabilizing the 3D structure of this regulatory domain. However, in the absence of a PAM sequence, we could not target it using CRISPR-CAS9 system and PAM-independent detection system will be required to address the function of this type of HLF binding motifs in the genome.

**Fig. 3.**
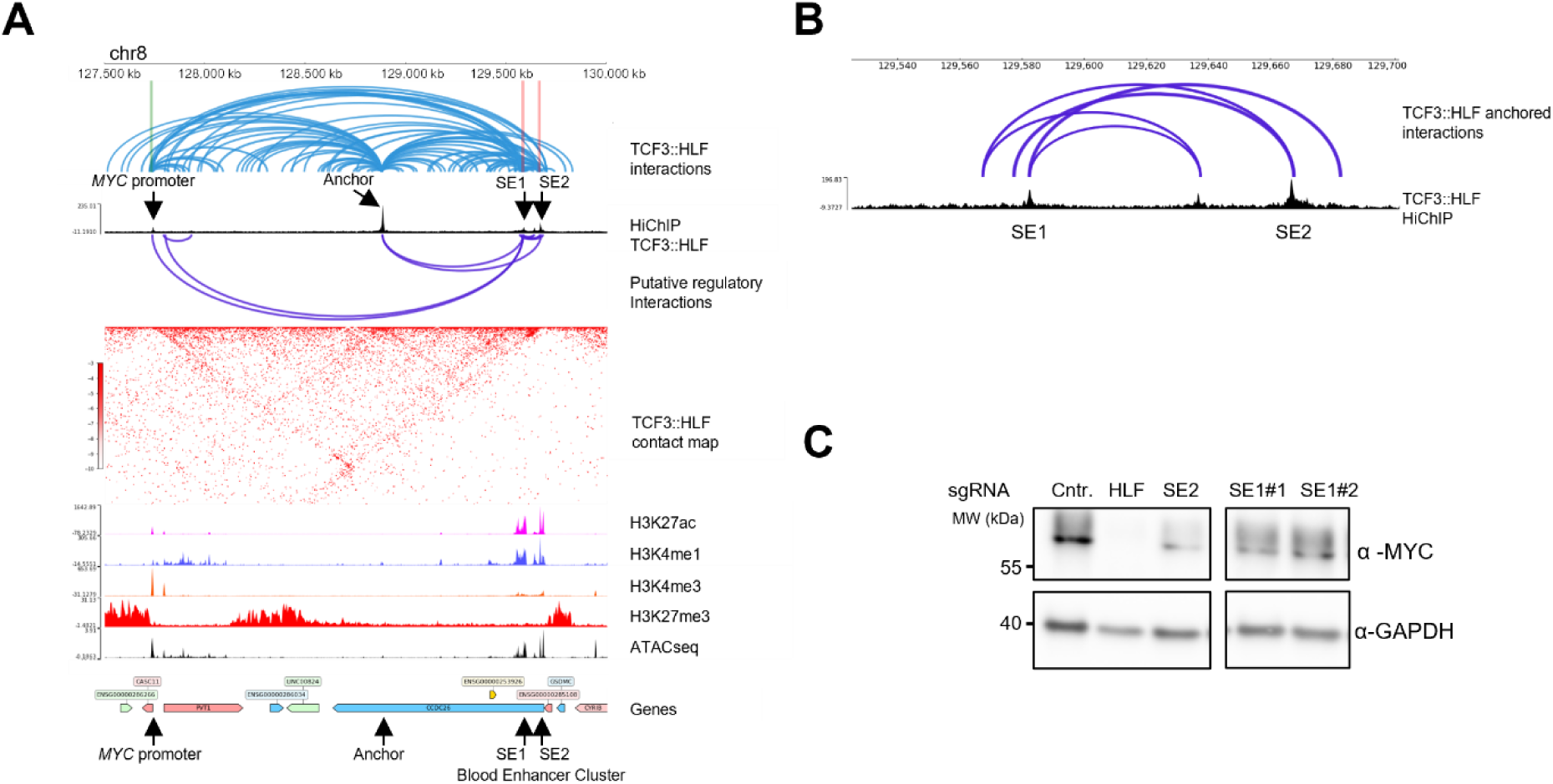
3D-genome interactions bound by TCF3::HLF at MYC promoter and BENC locus. (**A**) Visualization of TCF3::HLF genomic interactions at the MYC and Blood Enhancer Cluster (BENC) locus and the corresponding TCF3::HLF-HIChIP, H3K27ac, H3K4me1, H3K4me3, and H3K27me3 ChIPseq and ATACseq profiles. TCF3::HLF interactions identified by HiChIP are depicted as arcs. Interactions among regions with HLF motif are depicted with viola arcs. MYC promoter, anchor, SE1, and SE2 regions are indicated by arrows. (**B**) Magnification of TCF3::HLF-HIChIP profile at the BENC locus. TCF3::HLF interactions identified by HiChIP are depicted as arcs. SE1 and SE2 are indicated. (**C**) Western blot showing the expression of MYC in HAL-01 cells with sgRNA-control and sgRNAs targeting TCF3::HLF (HLF) or the HLF motif at SE2 or SE1 sequences. GAPDH serves as loading control.

Notably, we also found an additional binding site within the BENC locus, SE1, which interacts with SE2 and *MYC* promoter (**Fig. 3, A and B**). Both SE1 and SE2 were enriched for H3K27ac and H3K4me1, the chromatin signature of active enhancers. Disruption of the HLF site of SE1 by CRISPR was sufficient to reduce MYC expression similarly to cells with sgRNAs targeting SE2, demonstrating a role in enhancer function (**Fig. 3C**). These results indicated that TCF3::HLF-mediated *MYC* regulation requires TCF3::HLF association with both SE1 and SE2 sequences and the association with only one SE is not sufficient to drive MYC expression, implying a more complex regulation than initially thought. Together, these results confirm *MYC* as target of TCF3::HLF and illustrate the added value of 3D-genome information to study the TCF3::HLF-regulome.

### TCF3::HLF activation of MEF2C promotes expansion of leukemia cells

The *MEF2C* gene ranks among the most significant E-P interaction through TCF3::HLF binding (**Fig. 2F**, **Fig. 4A**). Moreover, *MEF2C* is significantly downregulated upon *TCF3::HLF*-KO in the parental HAL-01 cells, supporting that *MEF2C* is directly activated by TCF3::HLF. Remarkably, *MEF2C* is expressed homogeneously in most ALL subtypes, including TCF3::HLF and with the notable exception of MEF2D::BCL9 ALL, a rare ALL subtype with poor outcome (*51–53*) (**Fig. 4B**). MEF2C and MEF2D are evolutionarily conserved and important for the development of different tissues including blood (*54*). To determine the importance of MEF2C in TCF3::HLF positive B-ALL as a direct target of TCF3::HLF, we disrupted the TCF3::HLF binding site within the *MEF2C* enhancer by CRISPR. In TCF3::HLF positive B-ALL, disruption of the HLF binding site within the *MEF2C* enhancer markedly reduced MEF2C expression (**Fig. 4C**). ChIP analysis showed that TCF3::HLF association with the *MEF2C* enhancer was strongly reduced in cells treated with sgRNAs targeting the HLF motif compared to cells treated with sgRNA-control and sgRNA-3 targeting sequences downstream the HLF motif, indicating that HLF sequences are necessary for TCF3::HLF binding (**Fig. 4C**). Importantly, the deletion of this HLF motif caused the downregulation of MEF2C at RNA and protein levels, supporting a central role for this enhancer for MEF2C regulation in TCF3::HLF positive ALL (**Fig. 4D, fig. S3A)**.

**Fig. 4.**
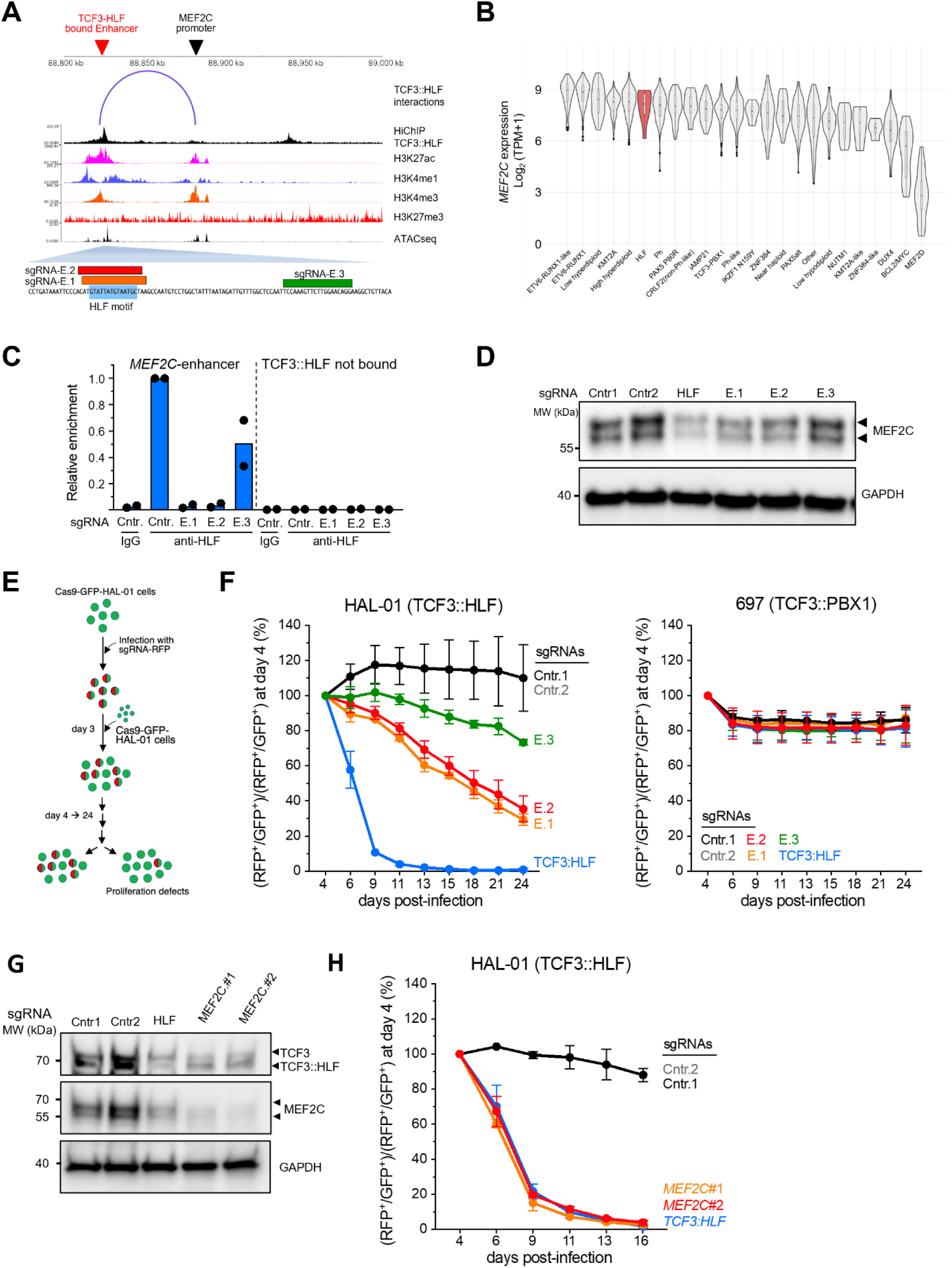
TCF3::HLF mediated regulation of MEF2C promotes expansion of leukemia cells. (**A**)Visualization of the genomic interactions between MEF2C promoter and its enhancer, bound by TCF3::HLF, including TCF3::HLF-HIChIP, H3K27ac, H3K4me1, H3K4me3, and H3K27me3 ChIPseq and ATACseq profiles. sgRNAs targeting the HLF motif at the *MEF2C*-enhancer (E.1, E.2) and downstream regions (E.3) are shown. (**B**) Violin plot of *MEF2C* expression across cohorts of different ALL subtypes. Datasets from the St. Jude Cloud (https://www.stjude.cloud) (46). (**C**) TCF3:HLF ChIP-qPCR for Cas9-expressing HAL-01 cells with sgRNAs targeting the *MEF2C-*enhancer HLF motif (E.1, E.2), downstream region (E.3) and sgRNA-Control (Cntr1,Cntr2). *MEF2C*-enhancer targeting primers and negative control primers targeting regions not bound by TCF3::HLF, were used. Values normalized to input and TCF3:HLF enrichment at *MEF2C*-enhancer in cells treated with sgRNA-Control (Cntr.). (**D**) Western blot of *MEF2C* expression in Cas9-expressing HAL-01 cells transduced with sgRNA-Control and sgRNAs targeting *TCF3::HLF* (HLF) or the *MEF2C*-enhancer (E.1, E.2, E.3). GAPDH serves as loading control. (**E**) Experimental scheme of the competitive assay: TCF3-rearranged cell lines (697, HAL-01), engineered to express Cas9-GFP, were transduced with RFP-co-expressing sgRNA vectors targeting TCF3::HLF (HLF), enhancer sites (E.1, E.2), or proximal region (E.3), and sgRNA-Control (Cntr1, Cntr2). **(F**) Competitive assay quantification in HAL-01 and 697 leukemia cells upon *MEF2C*-enhancer KO. Values correspond to the ratio between RFP+/GFP+ and GFP+ cell number at the indicated days relative to the ratio between RFP+/GFP+ and GFP+ cell number measured at day 4 post-transduction. (**G**) Western blot showing MEF2C and TCF3::HLF expression in Cas9-expressing HAL-01 cells transduced with control or targeting sgRNAs. GAPDH serves as a loading control. (**H**) Competitive assay quantification in HAL-01 and 697 leukemia cells upon *MEF2C*-KO. Values correspond to the ratio between RFP+ and GFP+ cell number at the indicated days relative to the ratio between RFP+ and GFP+ cell number measured at day 4 post-transduction. All results shown are mean values from two independent experiments.

To determine the functional consequences of the association of TCF3::HLF with *MEF2C* enhancer, we performed a cell competitive assay using two TCF3 rearranged cell lines (697 with t(1;19) and HAL-01 with t(17;19)) that were engineered for stable Cas9-GFP expression. The 697 ALL cell line harbors the TCF3::PBX1 fusion protein, which does not bind the HLF motif. Both cells were transduced with a lentiviral vector expressing red fluorescent protein (RFP) and sgRNAs targeting either *TCF3::HLF* or the TCF3::HLF-bound *MEF2C* enhancer site. Cell proliferation was quantified by flow cytometry by measuring the number of GFP^+^ and RFP^+^ cells from day 4 after infection until day 24. As expected, targeting *TCF3::HLF* in HAL-01 cells strongly impaired cell proliferation whereas it had no effect in 697 cells, indicating the specificity of the assay and further supporting the essential role of TCF3::HLF in t(17;19) positive ALL cells (**Fig. 4, E and F, fig. S3B**). Importantly, targeting the HLF site at the *MEF2C* enhancer also decreased cell proliferation in TCF3::HLF positive ALL but not in TCF3::PBX1 positive ALL cells, indicating an important and specific role of MEF2C in cell proliferation specifically for TCF3::HLF positive ALL. To further support the critical role of MEF2C in TCF3::HLF ALL cells, we performed the cell competitive assay in HAL-01 cells using sgRNAs targeting *MEF2C*. Western blot analyses confirmed MEF2C downregulation (**Fig. 4G**). Interestingly, reduction of MEF2C correlated with downregulation of TCF3:HLF, suggesting a regulatory loop between TCF3::HLF and MEF2C (**Fig. 4G**). Notably, *MEF2C*-KO impaired cell proliferation of TCF3::HLF in t(17;19) positive ALL cells at a rate similar upon *TCF3::HLF*-KO, underscoring the critical role of MEF2C (**Fig. 4H**). Collectively, these results indicated that the TCF3::HLF-mediated regulation of *MEF2C* through the interaction with the enhancer plays an important role in t(17;19) positive ALL.

### Activation of MEF2C by TCF3::HLF induces a gene expression program that maintains hematopoietic stem cell features and blocks lineage differentiation

To establish whether and how TCF3::HLF-mediated regulation of MEF2C can have an impact on gene expression programs linked to disease, we analyzed the RNAseq data of HAL-01 cells treated with sgRNA targeting the HLF motif of the *MEF2C* enhancer (*MEF2C*-enhancer KO). Cells were collected 13 days post-transduction, a time point at which HAL-01 exhibited approximately a 40% dropout rate in the competitive assay (**Fig. 4F**, **Fig. 5A**). We detected significant changes in gene expression (1021 upregulated and 969 downregulated genes, adj.*P* <0.05), including, as expected, the downregulation of *MEF2C* (**Fig. 5A**). Gene set enrichment analysis (GSEA) with HALLMARK signatures showed that the enhancer perturbation caused the downregulation of distinct gene expression programs, including *E2F* and *MYC* target genes (**Fig. S4A**), in agreement with results using independent models showing that MEF2C modulates *E2F* and *MYC* expression (*29*). Interestingly, GSEA with curated gene sets of biological pathways revealed positive enrichment of a B-cell progenitor signature after perturbation of the *MEF2C* enhancer, in addition to negative enrichment of genes involved in cell proliferation, pediatric cancers and stemness (**Fig. S4B**). GSEA utilizing a custom collection of published signatures from after *MEF2C*-KO or perturbed MEF2C activity (*29*, *55*, *56*) revealed overlapping signatures between genes downregulated upon *MEF2C*-enhancer KO and *TCF3::HLF*-KO and downregulated genes after MEF2C inactivation (**Fig. 5B**).

**Fig. 5.**
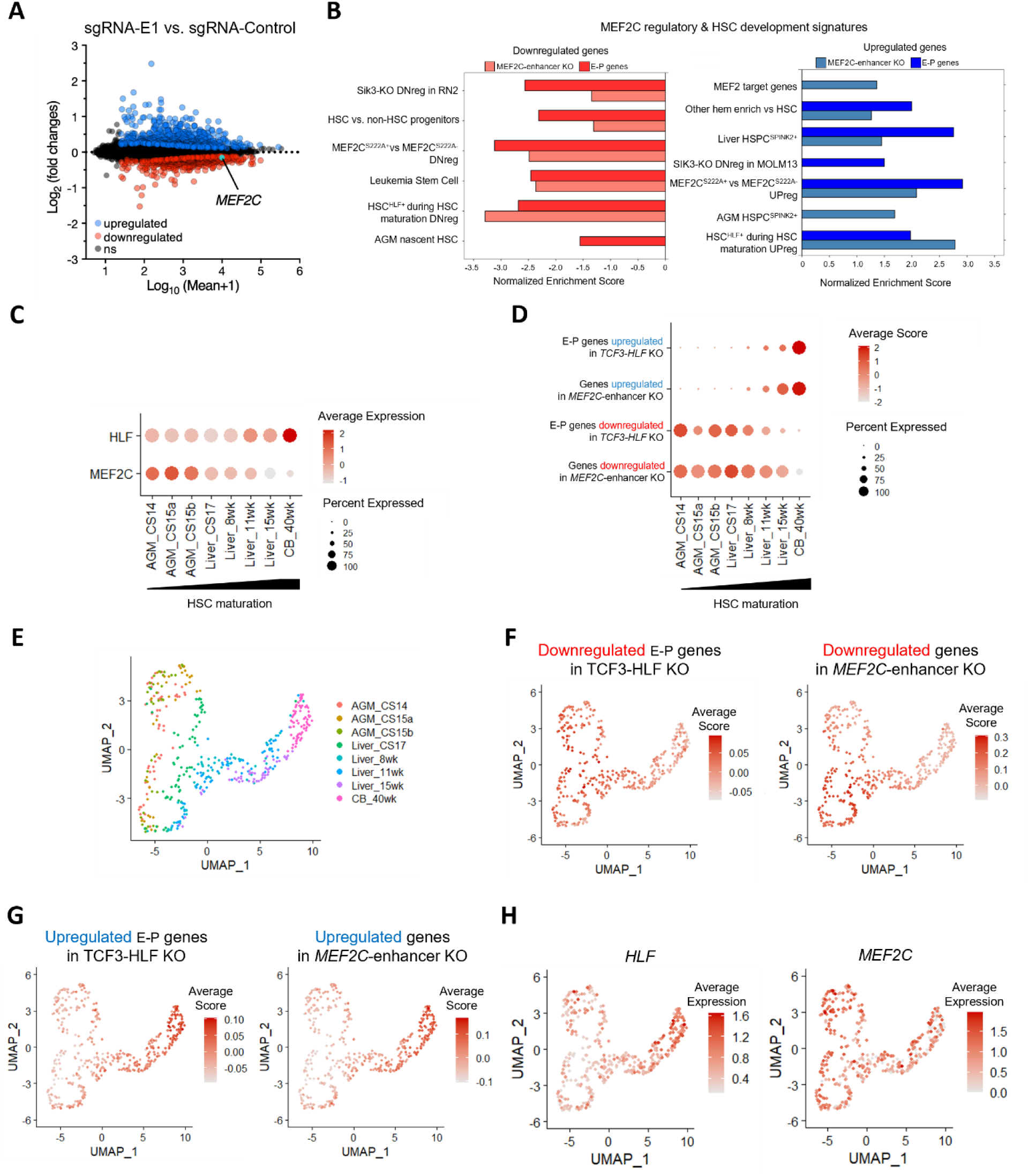
Activation of MEF2C by TCF3::HLF induces a gene expression program that maintains hematopoietic stem cell features and blocks lineage differentiation. (**A**) MA-plot showing differentially expressed genes upon MEF2C-enhancer KO in HAL-01 cells. Genes significantly upregulated (adj.p < 0.05) are represented in blue, while downregulated genes are in red. MEF2C gene is highlighted. (**B**) GSEA of upregulated and downregulated genes upon MEF2C-enhancer KO or *TCF3::HLF*-KO with MEF2C and HSC maturation gene sets from Calvanese et. al (Calvanese et. al. 2022). The x-axis displays the normalized enrichment score (NES) for significant values (adj.p <0.05). Right panel displays positive NES values and left panel shows negative NES values. (**C**) Dot plot comparing the expression of HLF and MEF2C during HSC maturation. Data are from (Calvanese et. al. 2022) (19). (**D**) Dot plot comparing module scores of top 200 upregulated and downregulated genes (adj.p <0.05) upon MEF2C-enhancer KO or of E-P genes that are up- and downregulated upon *TCF3::HLF*-KO in HAL-01 cells. Data are from (Calvanese et al. 2022). Carnegie stages (CS)14/15 aorta-gonad-mesonephros (AGM), CS17 to 15 weeks liver, 40 weeks cord blood (CB). The colour intensity and size of the dot plots correspond to average score and percent expression, respectively. (**E**) UMAP analysis showing the clustering of scRNAseq of HSC maturation. Data are from (Calvanese et al. 2022) (19). (**F, G**) HSC maturation scorecard dot plot showing top 200 upregulated (**F**) and downregulated genes (**G**) upon MEF2C-enhancer KO or from E-P genes differentially expressed upon *TCF3::HLF*-KO in HAL-01 cells. (**H**) UMAP analysis highlighting MEF2C and HLF expression.

Since HLF is one of the most specific TFs identifying human hematopoietic stem cells (HSC), we investigated gene sets of human HSC identity and maturation throughout development and aberrantly self-renewing progenitor-like cancer stem signature (*19*, *57*). *MEF2C-*enhancer KO expression profiles showed enrichment of signatures of both hematopoietic differentiation and HSC maturation (e.g., HSC^HLF+^ during HSC maturation UPreg) for upregulated gene targets, whereas signatures of undifferentiated HSC identity and immature nascent HSC were negatively enriched (e.g., HSC^HLF+^ during HSC maturation DNreg, HSC vs non-HSC progenitors). Leukemia stem cell signatures were also negatively enriched (**Fig. 5B**). These results suggest that MEF2C is a main driver of stemness programs in this TCF3::HLF positive leukemia. Furthermore, the same programs of HSC maturation and differentiation were also enriched in the HiChIP filtered dataset including TCF3::HLF-bound E-P loop genes that are regulated after *TCF3::HLF*-KO (**Fig. 5B**).

We further investigated how genes modulated upon *TCF3::HLF*-KO and *MEF2C*-enhancer KO are regulated throughout HSC maturation by interrogating publicly available single-cell RNAseq datasets mapping the transcriptomic changes in human embryonic HLF^+^ HSCs throughout maturation: from the aorta-gonad-mesonephros (AGM, Carnegie stage (CS) 14/15) region, the transition to the fetal liver until week 15, to cord blood (CB) (*19*). We found HLF expression increased with HSC maturation, whereas MEF2C expression was highest in the most immature embryonal subsets (AGM HSC) (**Fig. 5C**). Module score analysis of the differentially expressed genes after *TCF3::HLF*-KO with E-P contacts, as well as differentially expressed genes after *MEF2C*-enhancer KO, revealed a pattern concordant with the expression of MEF2C throughout maturation: upregulated genes are mostly expressed in mature HSC (i.e. 15 week fetal liver and CB) while downregulated genes are preferentially expressed in immature HSCs (i.e. AGM, early liver) (**Fig. 5, D to H**).

To determine how TCF3::HLF and MEF2C target genes are regulated throughout hematopoietic differentiation, we analyzed a published single-cell transcriptome map of HSCs and differentiated progenitors from human bone marrow (*58*). This data represents mature HSC, multipotent progenitors (MPP), megakaryocyte-erythroid progenitors (MEP), common myeloid progenitors (CMP) and granulocyte– monocyte progenitors (GMP), multi-lymphoid progenitors (MLP), Pre-B lymphocytes/natural killer cells (Pre-B/NK). *MEF2C* was expressed in lymphoid progenitor subsets (Pre-B/NK and MLP) compared to the more undifferentiated subsets, while HLF expression was almost exclusive to HSCs (**Fig. S4C**). Module score analysis revealed that E-P genes upregulated after *TCF3::HLF*-KO as well as upregulated genes after *MEF2C*-enhancer KO are normally expressed in lymphoid/B progenitors, consistent with signatures observed in TCF3::HLF leukemia cells (*20*). Downregulated genes were more expressed in less differentiated progenitors such as CMP and GMP (**fig. S4, D to H**). Collectively, these results suggest that TCF3::HLF and its regulation of *MEF2C* via enhancer binding contribute to driving programs associated with immature HSC and preventing the activation of lymphoid programs in leukemia.

## Discussion

In this study, we have delineated the role of the fusion protein TCF3::HLF in rewiring the 3D genomic landscape to promote leukemogenesis in t(17;19) positive ALL. Integration of HiChIP with RNAseq and histone marker ChIPseq data has revealed key enhancer-promoter interactions mediated by TCF3::HLF, providing a comprehensive view of the gene regulatory networks underlying this aggressive leukemia subtype.

This interaction map provides new insights into genes that are both positively and negatively regulated by the oncogenic fusion. Simultaneous direct repression by the oncogenic transcription factor must contribute to canalizing the abnormal cell fate, as suggested by many examples for fate decision through repression of genes relevant for differentiation (*59*). Our previous TCF3::HLF interactome analysis identified potential co-factors that can be recruited to alter chromatin states, such as members of SWI-SNF chromatin remodeling complex (SMARCA2, SMARCC1, and SMARCD2) and mediator factors promoting DNA loops interactions (YY1, CTCF, CGGBP1, and ZNF512) (*20*).

Specific E-P activity renders TCF3::HLF highly dependent on MYC. Besides sustaining metabolic states to favor proliferation, MYC can regulate chromatin decompaction in cells with high plasticity including activated B cells (*60*). We identify additional functionally relevant HLF binding sites within the blood enhancer cluster (BENC) that are critical determinants for leukemia propagation. HLF is established as one of the most specific HSC identity genes. HLF is expressed in naïve stems cells and multipotent hematopoietic progenitors (*17*, *18*, *61*) and in fetal HSCs (*19*, *62*, *63*). TCF3::HLF must reprogram a cell of origine to an immature lineage ambivalent state. Here we show that TCF3::HLF regulates *MEF2C* directly, in one of the most significant E-P loop detected by HiChIP in leukemia cells. MEF2C has been implicated as a regulator of the development in various tissues including of lymphoid T, B and NK lineages (*30*, *34*, *36*, *54*). MEF2C is activated as an oncogene by structural rearrangements in T-ALL (*64*), where it activates BCL2 (*36*). TCF3::HLF is also dependent on BCL2, similar to more immature lymphoid subtypes (*65*) and leukemic stem cells in acute myeloid leukemia (*66*). Here we identify the critical HLF binding site and enhancer that activates BCL2, which represents a relevant drug target in TCF3::HLF (*21*) that translated in molecular responses to venetoclax in combination with chemotherapy in compassionate treatment attempts and to the systematic recommendation for this leukemia subtype by the AIEOP-BFM Study Group in Europe (<u>NCT03643276</u>). Moreover, we show that perturbation of specific TCF3::HLF binding at the MEF2C enhancer did not only attenuate leukemia proliferation but also altered transcription to activate expression of genes associated with HSC maturation. Indeed, we detect MEF2C as a component of primitive hematopoietic stem cell gene expression signatures in fetal hematopoietic stem cells and adult stem and progenitor cells. The functional identification of MEF2C as a critical target of TCF3::HLF emphasizes the importance of specific E-P interactions in the maintenance of leukemic proliferation and stem-like characteristics.

Taken together, our results highlight fundamental components of the transcriptional circuitry that is rewired by this oncogenic fusion protein to drive the malignant phenotype. This reinforces the central need to develop new tools to target these fundamental driver mechanisms. This map of E-P interactions constitutes an important resource for future functional exploration of intragenic and extragenic TCF3::HLF mediated functions. A deep understanding of oncogenic programs will be required to pave the way for more effective combinations of targeted treatments for patients suffering from these devastating diseases.

## Material & Methods

### Cell lines and cell culture

The human leukemia cell lines harboring rearrangement variants of t(17;19) HAL-01 (*12*), and 697(DSMZ). All cell lines were grown in RPMI 1634 Media (GIBCO) supplemented with 10% fetal bovine serum (FBS), 1% glutamine (GIBCO), penicillin (100U/mL) and streptomycin (100 μg/mL) at 37°C and 5% CO2. Mycoplasma tests were routinely performed on all the cell lines.

### Western blotting

Whole cell lysates were prepared by resuspending cell pellets in Laemmli (BioRad#1610747) buffer with beta-mercaptoethanol and boiling for 6-10 min. Protein electrophoresis was performed using 4–20% Mini-PROTEAN® TGX™ Precast Protein Gels (BioRad) or 4–12% Criterion™ XT Bis-Tris Gel (BioRad). The proteins were transferred to Trans-Blot Turbo Midi 0.2 µm Nitrocellulose (BioRad) and immunoblotted using following antibodies; E2A (D2B1) Rabbit mAb #12258 (CellSignaling), c-Myc (D84C12) Rabbit mAb #5605 (CellSignaling), MEF2C (D80C1) Rabbit mAb #5030, GAPDH (D16H11) Rabbit mAb #5174T (CellSignaling), Anti-rabbit IgG-HRP Cat#A0545 (Sigma-Aldrich). Signal development was done with SuperSignal West Pico PLUS (ThermoFisher) ECL reagent in a Chemidoc Touch Imaging System (BioRad).

### Competitive Assays

Lentiviral transfections were performed as previously described (*20*). Cell lines stably expressing Cas9 and a GFP reporter previously established using the PX458 (Ran et al. 2013, Addgene: Cat#48138) plasmid backbone. Single guided RNAs were cloned into the shuttle plasmid co-expressing the RFP657 reporter (Huang et al. 2019, Addgene Cat#134968 ) as previously described. Efficiencies were determined by flow cytometry on day 3 after transduction. For the competitive assay co-transduced cells with Cas9-GFP and sgRNA-RFP were mixed 1:1 with Cas9-GFP mono-transduced cell lines. Starting from day 4 after transduction the distribution of the double (Cas9-GFP/sgRNA-RFP) and mono-(Cas9-GPF) transduced populations were analyzed by flow cytometry every 48 or 72 hours.

### ChIP and HiChIP sample processing

ChIP and HiChIP chromatin sample preparation followed the MNase HiChIP Dovetail® Kit (Cantata Bio) protocol v1.3 with modifications. In brief cells were harvested in batches of 30 million cells, pelleted and frozen at -80 degrees for at least 30min before fixation. Pellets were thawed on ice and washed once in PBS. Double cross-linking was performed using 1.7 mM EGS (ThermoScientific, #21656) in PBS for 45min at room temperature followed by 10 min in 1% formaldehyde (v/v) in PBS. Formaldehyde was quenched by adding glycine to a final concentration of 125 mM. Fixated cells were pelleted, snap frozen and stored at - 80 degrees until further use.

Fixed frozen cell pellets were allowed to thaw at room temperature and washed in 1x Dovetail® wash buffer according to protocol and resuspended in MNase digestion buffer using 100uL buffer per 10 million cells. 1000U of MNase was added per 100uL buffer and incubated for 6 min for chromatin digestion. The reaction was quenched by adding 50mM EGTA. Samples were diluted in MNase digestion buffer and supplemented with SDS to a final concentration of 0.1% and divided into two sonication vials 15 million cells in 400uL MNase digestion buffer each. Using maxed power settings eight cycles with 30 seconds ON and 30 seconds OFF, the chromatin was released from the cells (Micro Pulse Digenode). Chromatin fragmentation quality was validated with Agilent D5000HS ScreenTape System for DNA on a 2200 TapeStation platform, only digestions with a distribution of mononucleosomes between 40% and 70% were used for HiChIP.

For both ChIP and HiChIP 15ug of chromatin sample was used for pulldown. The chromatin was incubated over night with A/G magnetic Dynabeads coated with E2A (D2B1) Rabbit mAb #12258 (CellSignaling) or Rabbit IgG Isotype control (DA1E) #3900 (CellSignaling). Magnetic beads were washed with increasing stringencies of salt buffers with 150 mM NaCl, 500mM NaCl, 250mM LiCl and finally with TE buffer according to ChIP Abcam protocols, (Dovetail® washing procedure were not followed). For ChIP-qPCR, the samples were eluted at this stage by using the Dovetail® kit 1xReversal of crosslinks salt buffer and proteinase K incubated at 55 degrees for 15 min and 68 degrees for 45 min and purified using the PCR Purification kit from Qiagen.

For HiChIP the proximity ligation protocol was followed as instructed by the manufacturer. After streptavidin purification, the library preparation was done with the Dovetail Library prep module according to manufacturer’s protocol. The PCR amplification was done as recommended for the amount of pulled-down DNA material. Generated libraries were filtered by size using SpriSelect beads (Beckman Coulter). HAL-01 HiChIP biological replicates were pooled and sequenced using NovaSeq 6000 with 400M reads 2 x 150bp paired-end.

### HiChIP analysis

The HiChIP analysis pipeline documented in the HiChIP documentation release 0.1 by Dovetail® (https://hichip.readthedocs.io/en/latest/index.html) was followed with some modifications. HiChIP paired-end read files were trimmed with TrimGalore (v0.6.6) and filtered to eliminate short and low-quality reads (<50bp, phred 33), biological replicates were merged as recommended. HiChIP reads were aligned to the hg38 genome using Burrows Wheeler Aligner (*68*), using the bwa mem algorithm with settings as described in the HiChIP documentation release 0.1. Interaction events are extracted with the parse module of pairtools followed by sorting and splitting of the generated .pairsam file into .mapped.pairs and .mapped.bam files, the pairtools dedup step was omitted. We used mapped.bam files to generate primary aligned bed files for 1D MACS2 (*69*) peak calling (peaks, *n*=9552) as reference when calling genomic interactions with FitHiChIP(*70*), using 5kb bin size and 50kb to 3Mb range settings.

### Peak and motif analysis

To annotate the interactions where TCF3::HLF is directly involved as an anchor and identify the role of associated regulatory elements; we integrated our published histone ChIPseq data for HAL-01 in combination with HOMER v.4.7 (77) peakAnnotation (*71*), annotation of enhancer regions using both the ROSE output of MACS2 called broad peaks from H3K27ac ChIPseq data described above and further verified with Fantom5 Enhancer database. Binding sites with a HLF motif were identified with the aid of the Simple Enrichment Analysis (SEA) and Find Individual Motif Occurrences (FIMO) tools, from the MEME-suit (*72*), on MACS2 called peaks from TCF3::HLF HiChIP enrichment. Going by a 10Kb upstream and downstream cutoff window and integration of differential gene expression data from HAL-01 RNAseq after TCF3::HLF-KO, we defined putative promoter anchor sites associated with TCF3::HLF-bound anchors.

### ChIPseq and ATACseq data processing

The histone mark ChIPseq datasets for HAL-01 were downloaded from the European Nucleotide archive repository (ERP109232). Reads were trimmed using TrimGalore (v0.6.6) and aligned to the hg38 reference genome using bwa-mem (*68*). BigWig files were generated with deeptools (*73*). We selected regions of interest from HiChIP TCF3::HLF identified sites with HLF motif, and included top 75% of the regions with highest TCF3::HLF enrichment. Using deeptools computeMatrix analysis we proceeded to define the histone coverage using our own data (*20*) and publicly available ATACseq data, GSE186942 (*74*), at TCF3::HLF bound regions identified with HiChIP. Based on histone features we performed clustering with computeMatrix, into 3 main cluster groups.

### Calling enhancer clusters with ROSE

Enhancer clusters were called from H3K27ac ChIPseq data (*20*) as before using ROSE (*25*, *26*) on MACS2 broad peak calling outputs under the hg38 reference genome. Default parameters were chosen. In brief, H3K27ac peaks were first clustered on basis of gap distance, and enhancer clusters were identified by strong H3K27ac ChIPseq enrichment. Together with the Fantom5 database (*27*, *28*) annotations of TCF3::HLF-associated enhancer sites were made.

### RNAseq sample preparation

Total RNA samples were isolated using the Direct-zol™ RNA Miniprep with on column DNAse digestion, after lysis in TriReagent RT (MRC). RNAseq libraries were generated with the Truseq RNA Library Prep Kit for Illumina per manufacturer’s instructions and sequenced on a Novaseq 6000 with paired end 2 x 100bp.

### RNAseq expression and GSEA analysis

RNAseq reads were processed using Nextflow 23.04.4 nf-core/rnaseq 3.12.0 pipeline. In brief, sequenced reads were trimmed for adapter sequences/low-quality reads with trimmgalore 0.6.7 (parameters --phred33 --length 30 -j 4). Trimmed sequences were aligned to the hg38 genome (GCA_000001405.15_GRCh38_no_alt_analysis_set) using STAR alignment 2.7.9a. Read count quantification was done with Salmon 1.10.1. The same analysis pipeline was used to reanalyze the *TCF3::HLF*-KO dataset from the European Nucleotide archive under accession number ERP109232. Normalization and differential analysis were performed using R/Bioconductor version 4.3.1 software package Deseq2 in RStudio. Log foldchange (LFC) shrinkage was performed to generate more precis Log2 foldchange estimates using the ashr package for data used in MA-plot visualization (*75*). GSEA was done with the R package fgsea, the gene signatures used included HALLMARKS and curated gene sets of biological pathways (C2.signatures) (https://www.gsea-msigdb.org), and custom gene signature for MEF2C and HSC gene sets. We performed a stratification of the St Judes data by retrieving any genes with 3x higher expression median in TCF3::HLF positive leukemias than all other subtypes. Data are from the St. Jude Cloud datasets (https://www.stjude.cloud) (*46*).

### scRNAseq dataset comparisons

The expression of HLF and MEF2C were assessed in publicly available single cell RNAseq datasets from Calvanese et al. 2022 (human HSC maturation throughout development) and Pellin et al. 2018 (human bone marrow HSC and differentiated populations) using the R package Seurat (v3.1.2). Gene modules were compiled for the top 200 upregulated and downregulated genes (adj.p-value>0.05) after MEF2C enhancer KO or after TCF3::HLF-KO, selected for E-P genes and scores were calculated using AddModuleScore with default parameters. The module scores and the expression patterns of selected genes are shown using the DotPlot and FeaturePlot functions.

### Data visualization

RNAseq expression and GSEA analysis were visualized with plots generated using R/Bioconductor version 4.3.1 software package ggplot2 (https://ggplot2.tidyverse.org). HiChIP heatmaps were generated using HiC-Pro by merging valid pairs of replicates and processed as .hic files using juicer tools v1.22.01 function and visualized using Juicebox tools. Coolbox API with Jupetyr Notebook was used to summarize and generate genome visualizations for ChIPseq, ATACseq and HiChIPseq data. Circular plots were generated using R package circlize v0.4.16. Plots for PCR outputs and competitive assay were generated using GraphPad Prism version 10.2.0 for Windows, GraphPad Software, Boston, Massachusetts USA, www.graphpad.com.

### Statistical analysis

Experimental data are presented as means ± SD unless stated otherwise. Statistical significance was calculated by a two-tailed, t test with P < 0.05 considered statistically significant unless stated otherwise. Statistical significance levels are denoted as follows: *P < 0.05; **P < 0.01; ***P < 0.001 and ****P < 0.0001.

## Acknowledgement

We thank the Functional Genomics Center Zurich (FGCZ) of University of Zurich for the assistance in sequencing. Yun Huang for critical discussions and Q. Ngo for initial assistance with the computational analysis of pilot data.

## Funding

Comprehensive Cancer Center Zurich (CCCZ) fellow program (R.S and J-P.B)

Krebsliga Zürich (VP)

Krebsforschung Schweiz KFS-5298-02-2021 (J-PB)

European Research Council ERC-AdG-787074-NucleolusChromatin (RS).

## Author contributions

Conceptualization: VP, J-BP, RS

Methodology: VP, AD

Formal analysis: VP, BA, JA-G

Investigation: VP, AD

Visualization: VP, RS, AD, BA, JA-G

Supervision: VP, BB, J-PB, RS

Data Curation: VP

Writing—original draft: VP, RS

Writing—review & editing: VP, RS, J-PB, BB, AD, BG, JA-G, HKAM

Funding acquisition: VP, J-BP, RS

## Data and materials availability

HAL-01 *TCF3::HLF*-KO CRIPSR edited RNAseq datasets and HAL-01 Histone ChIPseq dataset from previous work are deposited in the European Nucleotide archive under accession number ERP109232. The ATACseq data for HAL-01 cells is under the GEO accession number GSE186942 (*74*).The single-cell transcriptomic data used to explore the expression of targets of interest during HSC development and differentiation are under the GEO accession number GSE162950 (*19*) and GSE117498 (*58*). All data are available in the main text or the supplementary materials.

## Supplementary Materials for

**Fig. S1.**
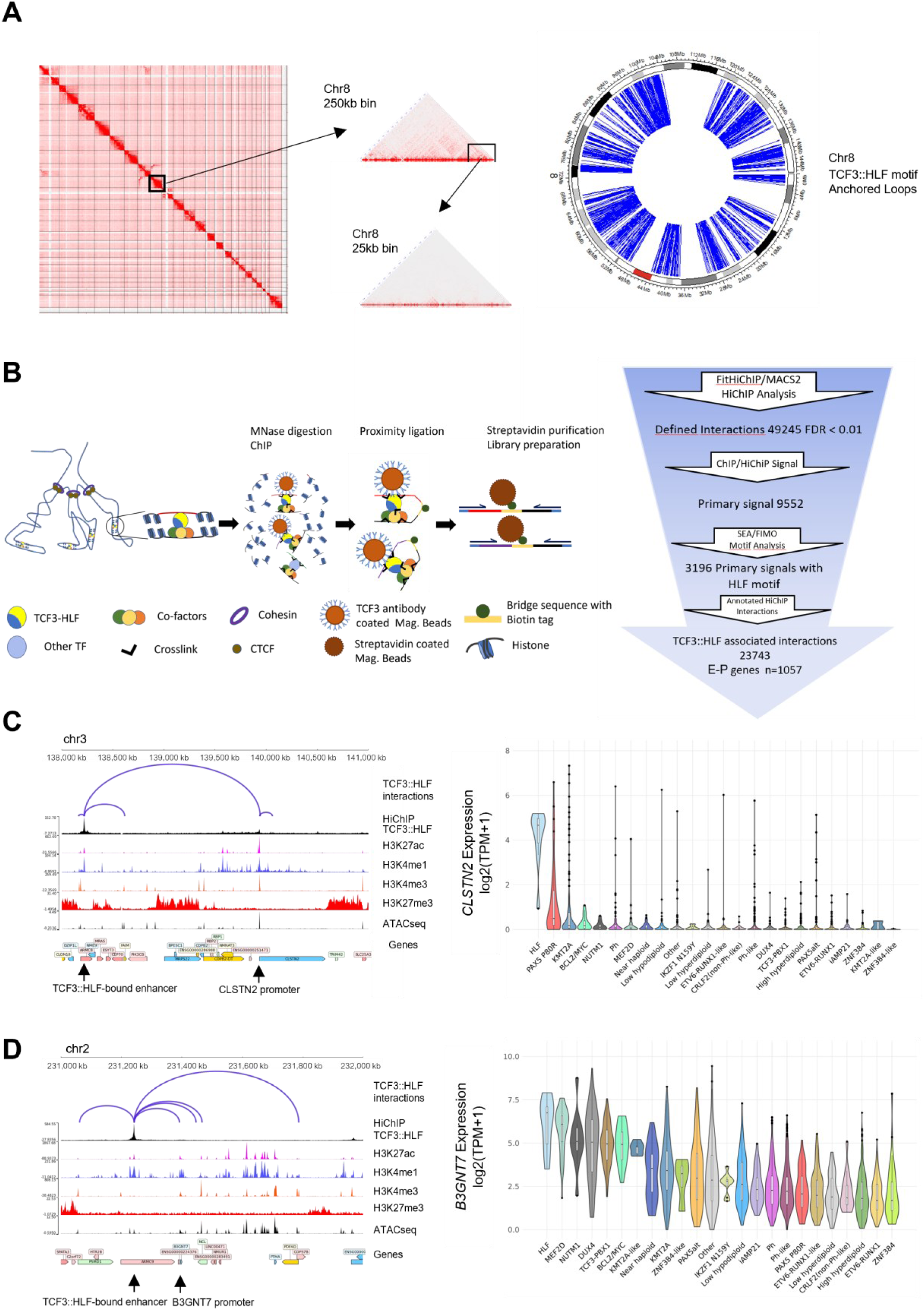
TCF3::HLF HiChIP pipeline. (**A**) HiChIP generated contact map of the TCF3::HLF mediated 3D genome landscape. Contact maps of chromosome 8 are shown at different resolutions, 250kB and 25kB bin sizes, revealing distribution of fusion protein associated interactions. Right panel. Circular ideogram of chromosome 8 showing the distribution of TCF3::HLF loops (blue arcs) with at least one HLF motif. (**B**) Schematic summary of the HiChIP strategy used to identify pairs of genomic interactions bound by TCF3::HLF and downstream analysis. (**C,D**) Left panels. Visualization of TCF3::HLF-associated interactions with *CLSTN2* (**C**) and *B3GNT7* (**D**) and the corresponding H3K27ac, H3K4me1, H3K4me3, and H3K27me3 ChIPseq and ATACseq profiles of HAL-01 cells. TCF3::HLF paired interactions identified by HiChIP are depicted as arcs. Right panel. Violin plots showing gene expression levels of *CLSTN2* (**C**) and *B3GNT7* (**D**) across different ALL subtypes. Data from the St. Jude Cloud (https://www.stjude.cloud) (*46*). HLF correspond to TCF3::HLF positive ALL.

**Fig. S2.**
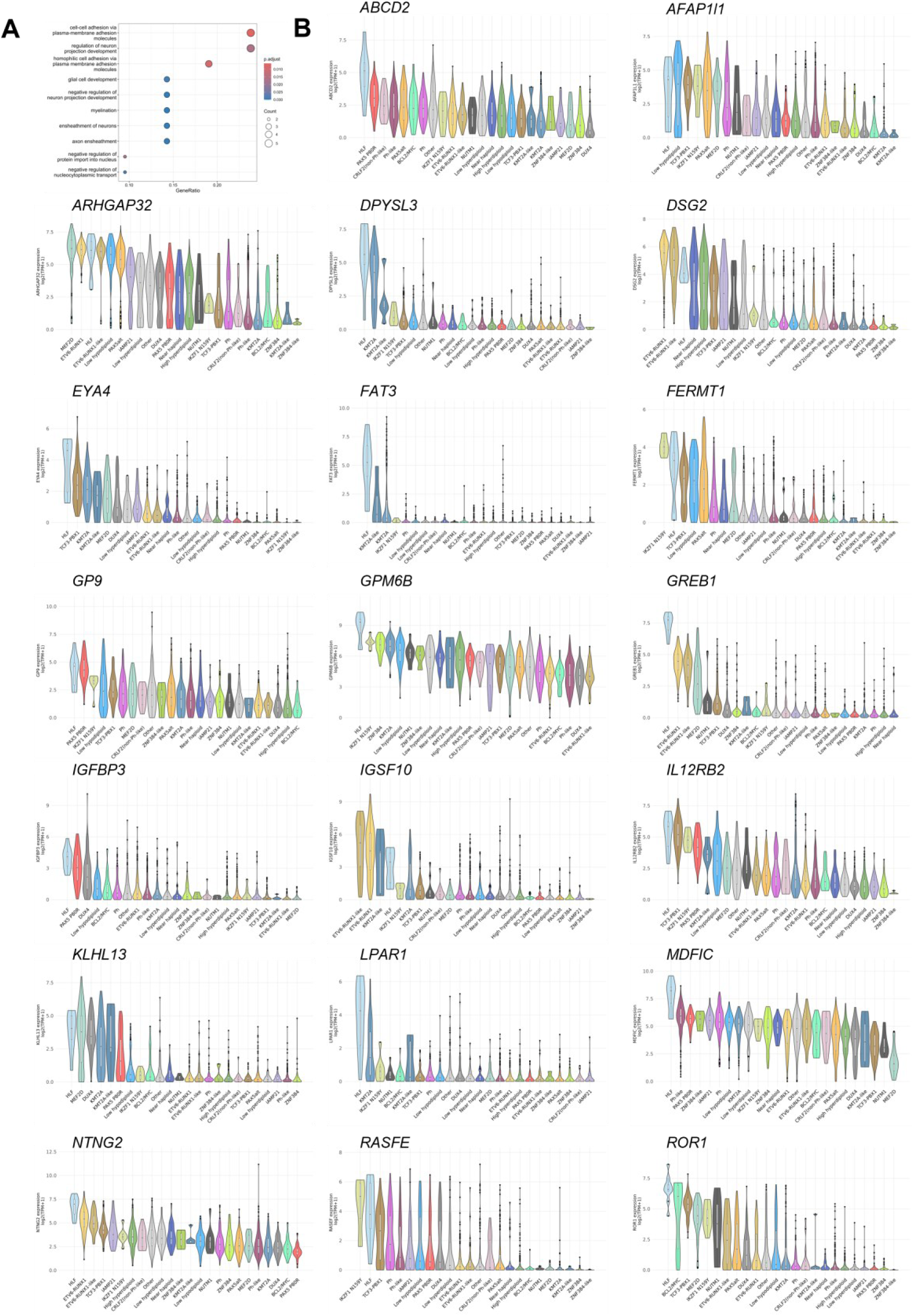
E-P gene expression among ALL subtypes. (**A**) Plots showing gene ontology terms for biological processes of E-P genes that are downregulated upon *TCF3::HLF*-KO and highest expression levels in TCF3::HLF (HLF) cohorts compared to other ALL subtypes. (**B**) Violin plots showing gene expression levels of 22 E-P genes across different ALL subtypes showing the highest mean expression in the HLF cohort compared to other ALL subtypes. Data are from the St. Jude Cloud (https://www.stjude.cloud) (46).

**Fig. S3.**
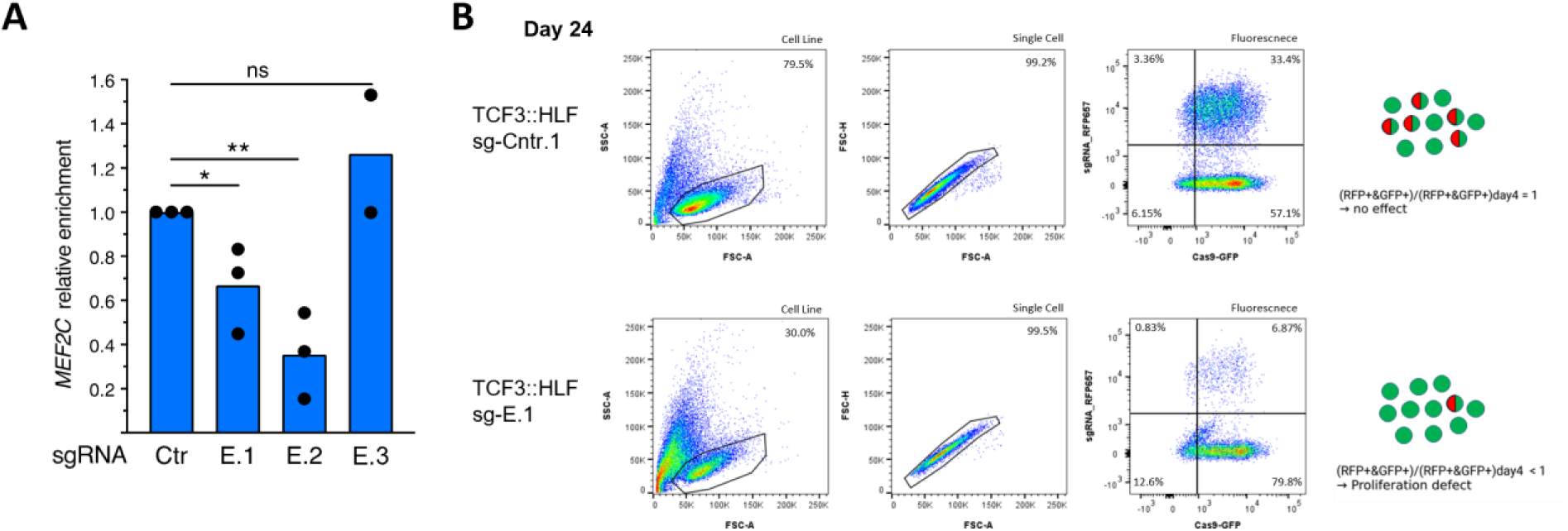
TCF3::HLF mediated regulation of MEF2C promotes expansion of leukemia cells. (**A**) RT-qPCR showing changes in *MEF2C* mRNA expression levels upon *MEF2C*-enhancer KO. Data were normalized to 18S and ACTB. Values are from three independent experiments. Statistical significance was calculated with unpaired T-test (* < 0.05; ** < 0.001; ns: not significant). (**B**) Scatterplot of competitive assay fluorescent output showing the comparative results at day 24 after transduction.

**Fig. S4.**
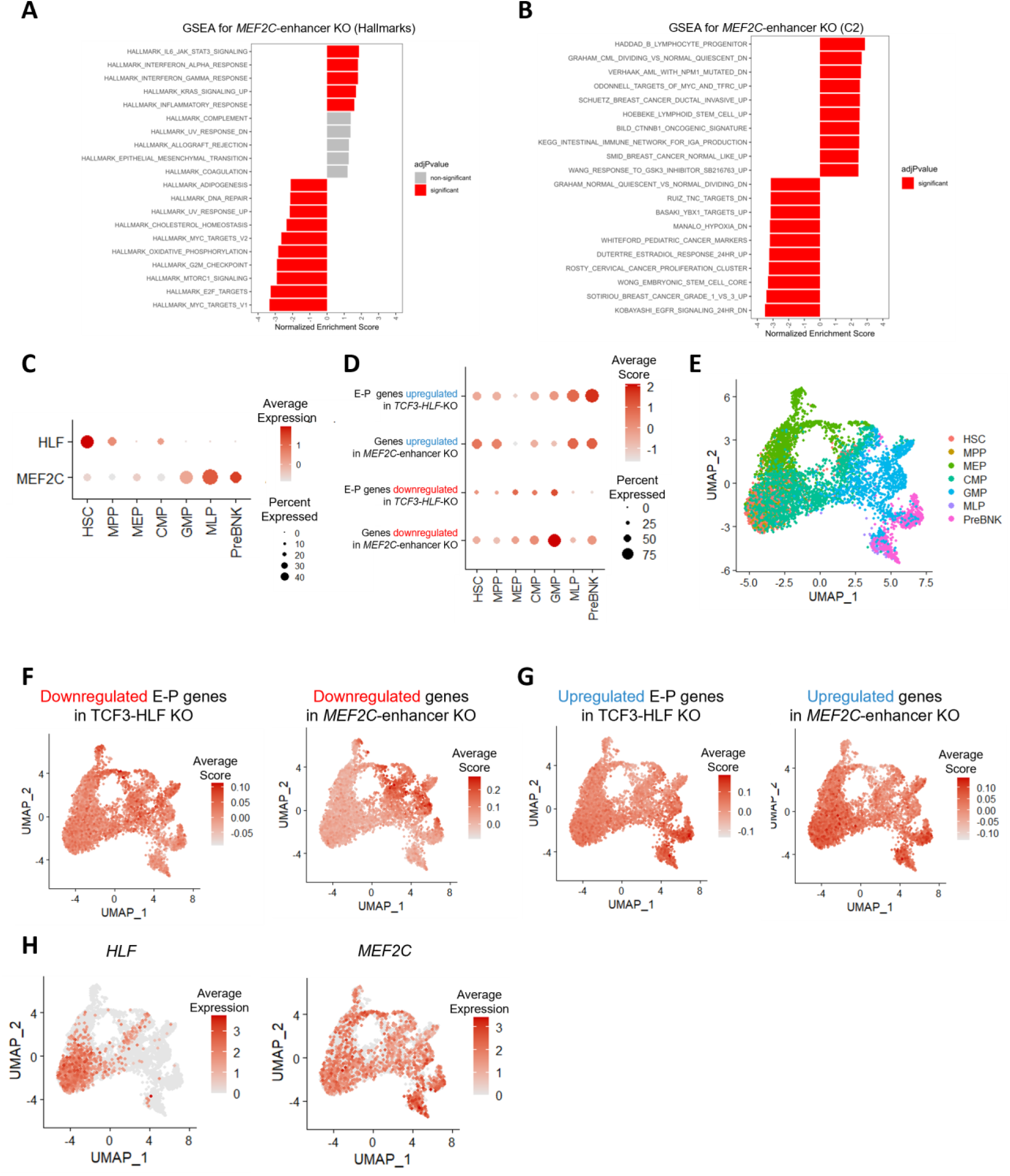
TCF3::HLF bound *MEF2C*-enhancer regulated gene signatures and expression of top genes in human bone marrow HSC and progenitors. (**A, B**) GSEA of upregulated and downregulated genes upon *MEF2C*-enhancer KO expression using HALLMARK and curated gene set from pathway databases and the biomedical literature (C2.signature) (https://www.gsea-msigdb.org). The x-axis displays the normalized enrichment score with significant values (adj.p <0.05). (**C**) Dot plot comparing the expression of *HLF* and *MEF2C* in the human bone marrow lymphoid, HSC and progenitor dataset by Pellin et al (*58*). (**D**) Dot plot comparing module score for top 200 upregulated and downregulated genes (adj.p >0.05) upon *MEF2C*-enhancer KO or E-P genes up- and downregulated upon *TCF3::HLF*-KO in HAL-01 cells. Using data from Pellin et al. (58). HSC, hematopoietic stem cells; MPP, multipotent progenitors; MLP, multi-lymphoid progenitors; Pre-B/NK, Pre-B lymphocytes/natural killer cells; MEP, megakaryocyte-erythroid progenitors; CMP, common myeloid progenitors; GMP, granulocyte–monocyte progenitors). The color intensity and size of the dot plots correspond to average score and percent expression, respectively. (**E**) UMAP analysis showing the clustering of scRNAseq of human bone marrow lymphoid, HSC and progenitor dataset. (**F,G**) Human bone marrow lymphoid, HSC and progenitor scorecard dot plot showing top 200 upregulated (**F**) and downregulated genes (**G**) upon MEF2C-enhancer KO or from E-P genes differentially expressed upon *TCF3::HLF*-KO in HAL-01 cells. (**H**) UMAP analysis highlighting MEF2C and HLF expression.

**Table S1.**
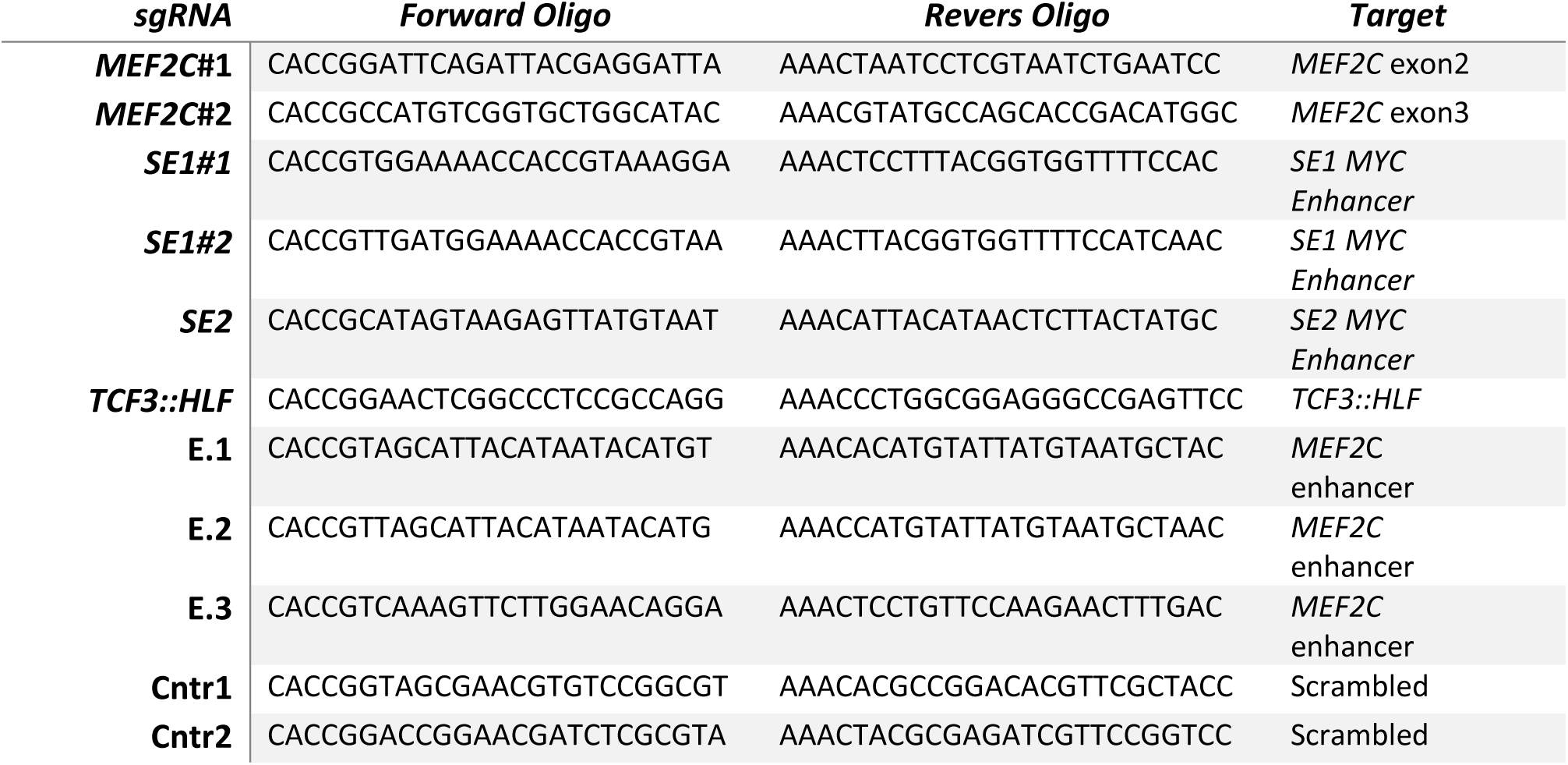
sgRNAs.

**Table S2.**
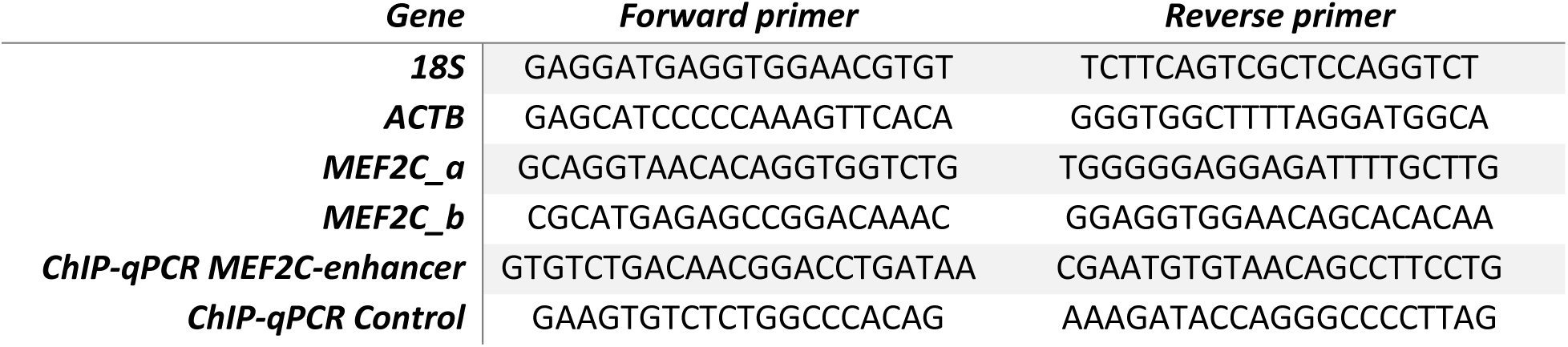
PCR primers.

